# Serotonin modulation of metabolism and stress response in *Pseudomonas fluorescens*

**DOI:** 10.1101/2025.04.03.646968

**Authors:** Barbora Waclawiková, Markus Schwalbe, Diana Ilyaskina, Semih Toptas, Nicola U Thome, Somayah S Elsayed, Anne de Jong, Gilles P van Wezel, Sahar El Aidy

**Author notes:** Corresponding author: Swammerdam Institute for Life Sciences, University of Amsterdam, Science Park 904, 1098 XH Amsterdam, Netherlands. Shared first author (Barbora Waclawiková, Markus Schwalbe).

## Abstract

**Background:** *Pseudomonas fluorescens* is a gram-negative bacterium with a remarkable metabolic and physiological versatility that enables it to adapt and colonize diverse ecological niches, including the human small intestine. While serotonin is primarily found in high concentrations in gut tissue, its levels in the lumen can be elevated in conditions such as celiac disease, where *P. fluorescens* is also found in increased abundance. The potential effects of serotonin on *P. fluorescens* in such contexts remain unclear.

**Results:** We demonstrate that *P. fluorescens* metabolizes serotonin primarily into 5-hydroxyindole-3-acetic acid (5-HIAA) and, to a lesser extent, into 5-hydroxytryptophol and N-acetylserotonin. Gene expression analysis revealed significant changes in oxidative stress-related pathways over time, and proteomic analysis confirmed the shifts seen particularly in amino acid catabolic pathways. Serotonin metabolism also enhanced bacterial resistance to oxidative stress, suggesting a protective role.

**Conclusions:** The findings reveal a novel mechanism by which serotonin modulates the metabolism and stress responses of *P. fluorescens*. This study provides insight into how *P. fluorescens* adapts to serotonin-rich environments, such as in celiac disease, and may inform future research on microbial interactions with host-derived metabolites in disease contexts.

## Introduction

*Pseudomonas fluorescens* is a gram-negative bacterium with a remarkable metabolic and physiological versatility, enabling it to adapt and colonize diverse ecological niches, including soil, water, plants and human digestive tract (Scales et al. 2014). *P. fluorescens* is a minor component of the gut microbiota in humans, where it mostly colonizes the small intestine (Ashorn et al. 2008; Dalwadi et al. 2001). *P. fluorescens* has been also linked to celiac disease (CeD), an autoimmune disorder primarily triggered by ingestion of gluten (Ashorn et al. 2008; Viitasalo et al. 2018; 2014; Petersen et al. 2019). Petersen, et al showed that gliadin reactive T cells involved in CeD pathogenesis cross-react with ubiquitous bacterial peptides, inferring microbial exposure as a potential environmental factor in CeD. The parental bacterial protein succinylglutamate desuccinylase (PFSGDS) derived from *P. fluorescens* mimics gliadin epitopes and activates gliadin reactive T cells derived from CeD patients (Petersen et al., 2019).

CeD has been also linked to the elevated levels of serotonin (up to 100 µM), due to the increased number of enterochromaffin cells (enterochromaffin cell hyperplasia) (Coleman et al. 2006). However, whether there is a link between *P. fluorescens*, and elevated levels of serotonin, remains elusive.

In the gut, serotonin is released either basolateral into the lamina propria, or apically into the lumen (Bertrand and Bertrand 2010), suggesting that gut microbes are exposed to the host-derived serotonin. Indeed, gut microbes, in particular, a subset of its spore-forming members, have been shown to take up luminal serotonin via a homologue of the mammalian serotonin selective reuptake transporter (Fung et al. 2019). Furthermore, serotonin can be sensed by the gut microbiota by a sensor kinase CpxA, a bacterial membrane-bound histidine sensor kinase (Kumar et al. 2020). Serotonin has also been demonstrated to act as a regulator of the gut bacterial virulence and the bacterial growth, especially in *Escherichia coli* and *P. fluorescens* (Kumar et al. 2020; Biaggini et al. 2015; Oleskin et al. 1998). Here, we explore the dynamic effects of elevated serotonin levels on *P. fluorescens* using RNA-seq and proteomics to uncover changes in gene expression and protein profiles. We also demonstrate that *P. fluorescens* metabolizes serotonin and how this process influences bacterial growth and key molecular pathways.

## Results

### *P. fluorescens* metabolizes serotonin into multiple metabolic products, similar to human end products

To investigate whether gut microbiota can metabolize serotonin, we screened 14 aerobic and 18 anaerobic bacterial strains from diverse genera, including, *Pseudomonas*, *Escherichia*, *Bacillus*, *Corynebacterium*, *Enterococcus*, *Fusobacterium*, *Desulfovibrio*, *Acidaminococcus*, *Veillonella*, *Clostridium*, *Bacteroides*, *Atopobium*, *Eggerthella*, *Dialister*, *Ruminococcus* and *Prevotella* (**Supplementary Table 1**). Notably, only strains within the *Pseudomonas* genus, particularly, *P. fluorescens* strains, were capable of metabolizing serotonin. Among the tested *P. fluorescens* strains, *P. fluorescens* MFY63, a human isolate (Chapalain et al. 2008), exhibited the most efficient conversion of serotonin (**Supplementary Table 1**), and was therefore selected for further study.

To investigate serotonin catabolism in *P. fluorescens*, the strain was incubated aerobically, being a strict aerobe, with 100 µM serotonin at 37 °C for 24 hours. Cultures were extracted using ethyl acetate and analyzed via Liquid Chromatography-Tandem Mass Spectrometry (LC-MS/MS). Data processing with MZmine (version 2.53, (Pluskal et al. 2010)) and comparative metabolomics analysis using MetaboAnalyst (Xia et al. 2009) identified eight mass features significantly enriched in serotonin-treated *P. fluorescens* cultures (**Figure 1A,B**). Two mass features were excluded: one (*m/z* 329.1395, [M+H]^+^) was also present in the control medium with serotonin, and the other (*m/z* 437.2183, [M+H]^+^) was a dimeric ion of another feature (**Figure 1A**). The remaining upregulated mass features were dereplicated by comparing their high-resolution MS and MS/MS spectra against chemical databases such as ChemSpider and Reaxys. These features included: (**1**) *N*-acetylserotonin, (**2**) 2-oxo-*N*-acetylserotonin, (**3**) 5hydroxy-3-vinyl-1*H*-indole, (**4**) *N*-(4-amino-1-hydroxybutyl)-serotonin, (**5**) derivative of hydroxy-vinyl-indole, and (**6**) unidentified metabolite (**Figure 1B**). The identity of N-acetylserotonin was confirmed by matching the observed peak with a commercially available standard (**Figure 1C**). In humans and other eukaryotic organisms, serotonin is degraded via multiple pathways, producing various metabolic products (Roth et al. 2021; Squires et al. 2010; Fanciulli et al. 2020). We hypothesized that *P. fluorescens* may similarly metabolize serotonin into several compounds, including: 5-hydroxyindole-3-acetic acid (5-HIAA), 5-hydroxytryptophol, 5-methoxytryptamine, formyl-5-hydroxy-kynureamine and *N*-methylserotonin. To test this, we specifically searched the LC-MS/MS data for these metabolites. Minor peaks corresponding to 5-HIAA (**Figure 1D**) and 5-hydroxytryptophol (**Figure 1E**) were detected exclusively in serotonin-treated cultures. However, these peaks were not identified in the general metabolomics workflow due to their low signal intensity, which was close to the noise level.

**Figure 1.**
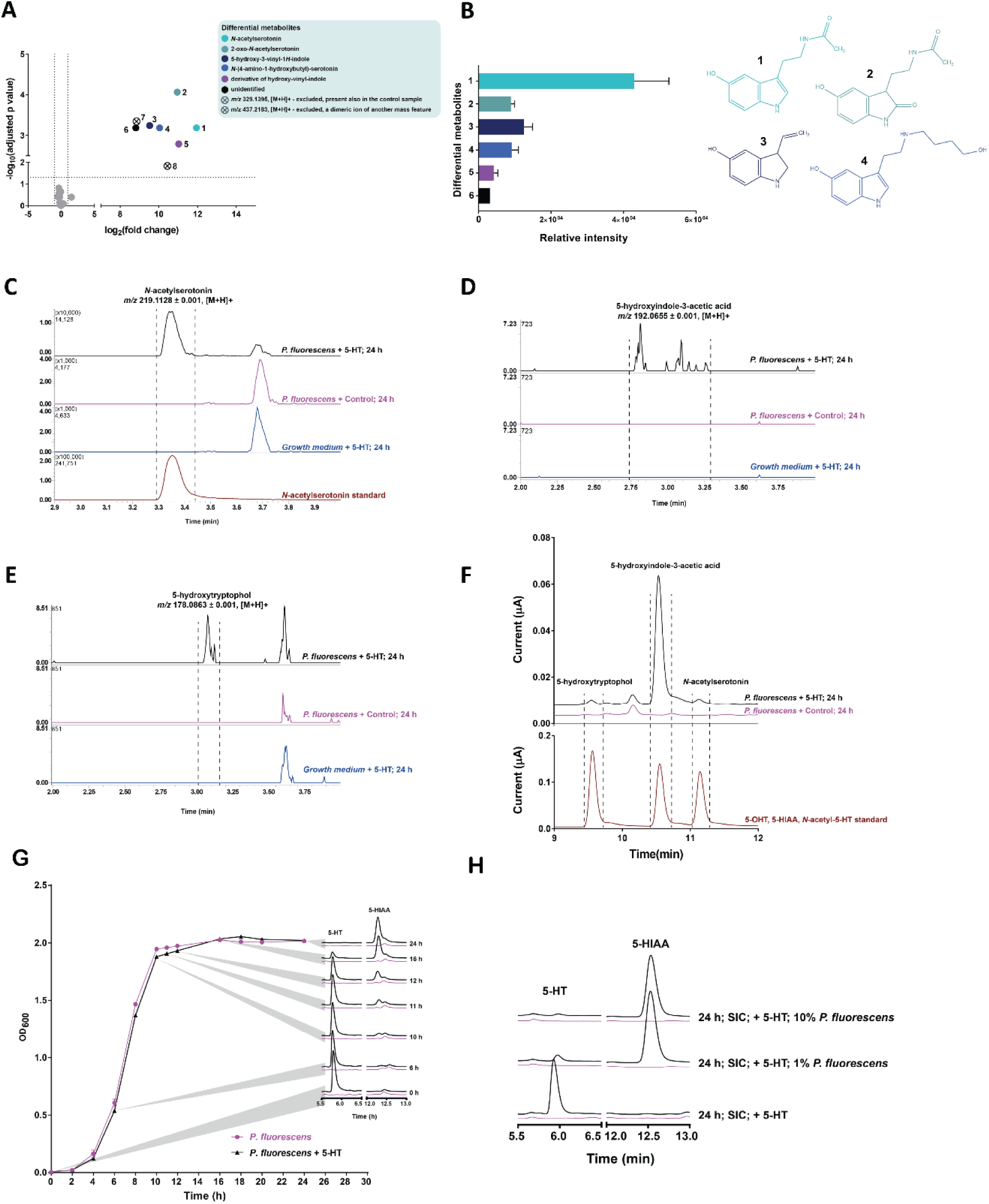
Serotonin is metabolized by *P. fluorescens* to numerous metabolic products, similar to the human metabolic end products. (A) Volcano plot of differential metabolites significantly abundant in the extracts of *P. fluorescens* cultures treated versus untreated with serotonin. Identification level is 1 for *N-*acetylserotonin and 2 for the rest (Sumner et al. 2007). (B) Bar graph (metabolite 1 – 6) and corresponding chemical structures (metabolite 1 – 4) of differential metabolites significantly abundant in *P. fluorescens* treated with serotonin. (**C – F**) LC-MS/MS chromatograms show the formation of *N-*acetylserotonin (**C**), 5-hydroxyindole-3-acetic acid (**D**) and 5-hydroxytryptophol (**E**) in *P. fluorescens* after incubation for 24 h (black line) and compared to the *P. fluorescens* after incubation for 24 h with control (purple line), growth medium with added serotonin only (blue line). Brown line in **C** indicates *N*-acetylserotonin standard. Filtering conditions (*m/z* range) were *m/z* 219.1128 ± 0.001 for *N*-acetylserotonin, *m/z* 192.0655 ± 0.001 (*m/z* 192.0645 – 192.0665) for 5-hydroxyindole-3-acetic acid and *m/z* 178.0863 ± 0.001 for 5-hydroxytryptophol. (**F**) HPLC chromatograms show the ratio of 5-hydroxytryptophol, 5-hydroxyindole-3-acetic acid and *N-*acetylserotonin. **(G)** Growth curve of *P. fluorescens* stimulated with serotonin (black line) or control (purple line). Grey highlights indicate when samples for HPLC-ED analysis were collected and point to the chromatograms of serotonin degradation over time in *P. fluorescens* incubated at 37 °C in EBB with agitation with serotonin (black line) or control (purple line) for 0, 6, 10, 11, 12, 16 and 24 h. Chromatograms represent one example of 3 biological replicates. Curve represents one example of 4 biological replicates. **(H)** Overnight culture of rat SIC cultured at 37 °C with agitation with serotonin for 24 h, in the absence or presence of 1 % or 10 % of *P. fluorescens*. Chromatograms represent one example of 5 biological replicates. Abbreviations: 5-HT, serotonin; 5-HIAA, 5-hydroxyindole-3-acetic acid; SIC, small intestinal content.

To determine the relative proportions of these metabolites in P. fluorescens after 24 hours of serotonin exposure, we analyzed the samples using High-Performance Liquid Chromatography coupled with electrochemical detection (HPLC-ED) (**Figure 1F**). The results revealed that 5-hydroxytryptophol, 5-HIAA, and N-acetylserotonin were produced in a ratio of 2.6:94.2:3.2, indicating that 5-HIAA is the predominant metabolic product of serotonin in *P. fluorescens*.

Next, to determine the growth phase during which *P. fluorescens* metabolizes serotonin to 5-HIAA, the bacterium was cultured in Enriched Beef Broth (EBB) supplemented with 2 g/L glucose or in minimal medium (M9). Growth was monitored in the presence and absence of serotonin for 24 hours, with samples collected at different phases: early exponential (6 h), early stationary (10, 11, 12 h), and late stationary (16, 24 h) for HPLC-ED analysis (**Figure 1G**). The results showed that serotonin degradation began after 10 hours of growth, with complete metabolism to 5-HIAA occurring after 24 hours (**Figure 1G**). Notably, serotonin had no impact on the growth of *P. fluorescens* under these conditions (**Figure 1G**).

To assess whether *P. fluorescens* can degrade serotonin in the presence of other small intestinal bacteria, we used rat proximal small intestinal content (SIC) to simulate the gut environment. SIC was cultured aerobically with serotonin and analyzed by HPLC-ED. The results showed that serotonin was metabolized to 5-HIAA within 24 hours only when *P. fluorescens* (at 1% or 10% inoculation) was present (**Figure 1H**). In contrast, SIC incubated with serotonin alone did not exhibit serotonin degradation after 24 hours (**Figure 1H**). Taken together, our results show that, similar to humans and other eukaryotic cells, *P. fluorescens* metabolizes serotonin into multiple products, including 5-hydroxytryptophol, 5-HIAA, and N-acetylserotonin, with 5-HIAA being the most abundant product. This raises the question of whether serotonin metabolism induces molecular modifications in *P. fluorescens*.

### Serotonin elicits metabolic shifts in *P. fluorescens*

Due to the unavailability of the *P. fluorescens* MFY63 genome, we performed whole genome sequencing and subsequent *de novo* genome assembly, which yielded 66 contigs, with a mean C+G content of 63.64% and a total genome size of 7.4 Mbp. We identified 6,768 coding sequences, which were annotated using the Prokka pipeline (Seemann 2014).

To investigate whether exposure to serotonin induces specific molecular modifications in *P. fluorescens*, RNA-seq was performed on samples collected at various time points following treatment with 100 µM serotonin (**Figure 2A**). Global gene expression analysis of the RNA-seq data from serotonin-treated samples versus controls was conducted using the Trex-2 analysis pipeline, a parameter-free statistical tool for RNA-seq gene expression (de Jong et al. 2015). The analysis revealed significant changes in gene expression, with 11 genes showing differential expression at the 10-hour time point and 93 genes at the 12-hour time point (**Figure 2B**; **Supplementary Table 2 – 3;** false discovery rate (FDR) ≤ 0.05, fold change ≥ 2), respectively. No significant gene expression changes were observed at 6, 8, or 24 hours based on the selected cutoffs (FDR ≤ 0.05, fold change ≥ 2).

**Figure 2.**
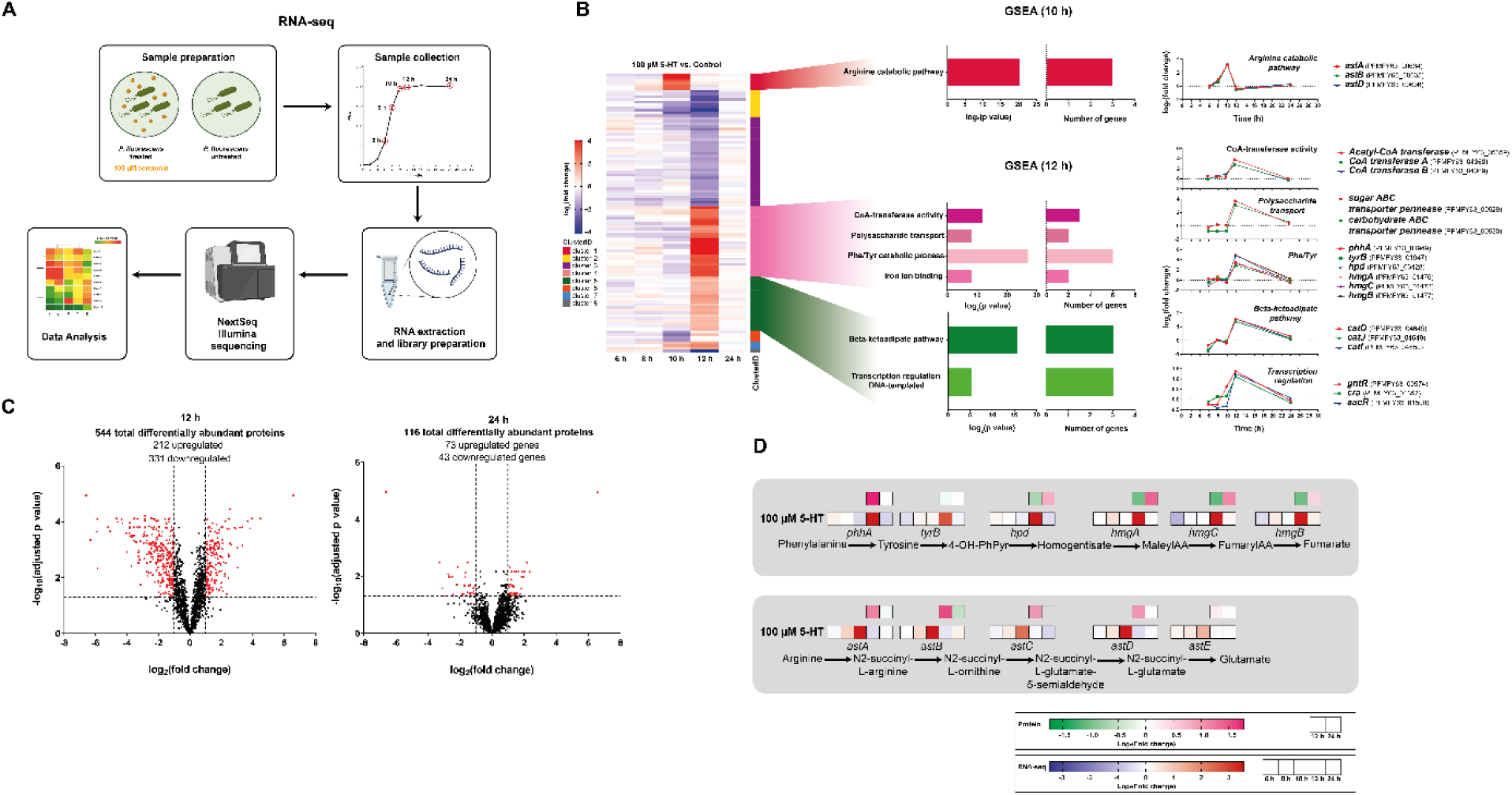
Serotonin metabolism protects *P. fluorescens* from oxidative stress possibly via upregulation of phenylalanine and tyrosine catabolic pathways. (A) RNA-seq experimental design. (B) Heat map of the genes that are significantly regulated (at 10 and 12 h) in the presence of serotonin as compared to the untreated control. Data represent log_2_(fold change). FDR ≤ 0.05, fold change ≥ 2. GSEA analysis of upregulated genes from stimulation with serotonin (FDR ≤ 0.05, fold change ≥ 2) after 10 h and 12 h. Data show −log_2_(p value) for specific GO biological pathways (bar graphs on the left); number of genes involved in the regulated pathways (bar graphs in the middle); and, expression patterns of individual genes involved in the regulated pathways (graphs on the right). (C) Volcano plots of differentially abundant proteins after 12 h (left panel) and 24 h (right panel) of stimulation with serotonin in *P. fluorescens*. (D) Overview of phenylalanine, tyrosine and arginine catabolic pathway and its regulation by serotonin on the RNA and protein level.

Next, Gene Ontology (GO) analysis was performed to identify biological pathways affected by serotonin treatment, using the FUNctional Analysis and Gene Set Enrichment Analysis for Prokaryotes (FUNAGE-Pro) (de Jong, Kuipers, and Kok 2022). The data revealed that a total of 1 and 6 Gene Ontology (GO) biological terms were overrepresented in *P. fluorescens* exposed to 100 µM serotonin at the 10-hour and 12-hour time points (early stationary phase), respectively (**Supplementary Table 4**). The GO pathways involved at the 10-hour time point included the arginine catabolic pathway, while those at the 12-hour time point included CoA-transferase activity, polysaccharide transport, phenylalanine and tyrosine catabolic pathways, iron ion binding, beta-ketoadipate pathway, and transcription regulation (**Figure 2B**).

To further explore the functional impact of serotonin on *P. fluorescens*, we performed proteomic analysis on samples collected at the 12-hour and 24-hour time points. These time points were selected based on RNA-seq results, where significant differential regulation of genes was observed at 12 hours. We hypothesized that the later time point (i.e., 24 hours) would show persistent changes at the proteomic level that were not detectable at the RNA level (**Figure 2B**). Proteomic data analysis using MSstats, an R package for statistical analysis of quantitative mass spectrometry-based proteomic experiments (Choi et al. 2014) identified 544 differentially abundant proteins at the 12-hour time point (**Figure 2C, Supplementary Table 5;** FDR ≤ 0.05, fold change ≥ 2) and 116 differentially abundant proteins at the 24-hour time point (**Figure 2C, Supplementary Table 6;** FDR ≤ 0.05, fold change ≥ 2). Further analysis of the proteomic data using FUNAGE-Pro revealed numerous biological pathways significantly overrepresented in both upregulated and downregulated gene sets at both time points (**Supplementary Table 7**).

Notably, amino acid catabolic processes, particularly the phenylalanine, tyrosine, and arginine catabolic pathways, were prominently upregulated at both the RNA and protein levels (**Figure 2D**). The phenylalanine and tyrosine catabolic processes were the only complete pathways, consisting of 6 genes (*phhA*, *tyrB*, *hpd*, *hmgA*, *hmgC*, *hmgB*), whose transcription was significantly upregulated at the 12-hour time point following serotonin stimulation. At the 24-hour time point, 2 out of these 6 proteins were significantly more abundant in the serotonin-treated *P. fluorescens* (**Figure 2D**). Similarly, the arginine catabolic process showed upregulation at both the RNA level (3 out of 5 genes) after 10 hours of serotonin stimulation and at the protein level (4 out of 5 proteins) at the 12-hour time point (**Figure 2D**). Of note in our proteome data was the minor upregulation (FDR = 0.04, log2 fold change = 0.3) of a homologue of succinylglutamate desuccinylase (PFSGDS, PFMFY63_00638) compared to the control at the 12-hour time point. This enzyme and its derived peptides have been reported to mimic gluten peptides when displayed by antigen-presenting cells, thereby activating T cells in patients with celiac disease in the same manner as gluten (Petersen et al. 2019).

Taken together, the de novo sequencing of *P. fluorescens* MFY63 genome enabled a comprehensive exploration of the transcriptomic and proteomic responses upon serotonin exposure. The combined RNA-seq and proteomics data reveal that serotonin metabolism induces significant upregulation of various functional pathways in *P. fluorescens*, with a particular focus on amino acid catabolic pathways, including phenylalanine, tyrosine, and arginine catabolism. These findings provide new insights into the molecular adaptations of *P. fluorescens* in response to serotonin, highlighting its potential role in modulating microbial metabolic networks.

### Serotonin metabolism protects *P. fluorescens* from oxidative stress

Serotonin has been previously described to possess antioxidant properties (Kang et al. 2009; Wan et al. 2018; Azouzi et al. 2017; Gonçalves et al. 2022; Liang et al. 2020). Similarly, the phenylalanine, tyrosine, and arginine degradation pathways, which were significantly overrepresented in our RNA-seq and proteomic analyses (**Figure 2B,D**), are known to play a role in bacterial responses to oxidative stress (Id et al. 2019; Tiwari et al. 2018; Ipson and Fisher 2016). Since *P. fluorescens* was grown under aerobic conditions in the initial serotonin catabolism study, it was exposed to oxidizing agents produced during aerobic respiration (da Cruz Nizer et al. 2021). This led us to hypothesize that serotonin metabolism may provide a protective effect against oxidative stress in *P. fluorescens.* To test this hypothesis, *P. fluorescens* was cultured in the presence or absence of serotonin and exposed to different concentrations of hydrogen peroxide (H₂O₂) (40 µM and 80 µM) to induce oxidative stress, as previously reported (Sies 2017). The bacterial growth (OD_600_) was monitored spectrophotometrically over a 48-hour period. The results demonstrated that serotonin significantly increased the carrying capacity (k), which reflects the maximum OD_600_, in the presence of both H₂O₂ concentrations (**Figure 3A,B)** compared to the control. Notably, serotonin did not induce any changes in the carrying capacity in the absence of H_2_O_2_ (**Figure 3A,B**). Collectively, the data suggest that serotonin acts as an antioxidant, protecting *P. fluorescens* from oxidative stress induced by H_2_O_2_.

**Figure 3.**
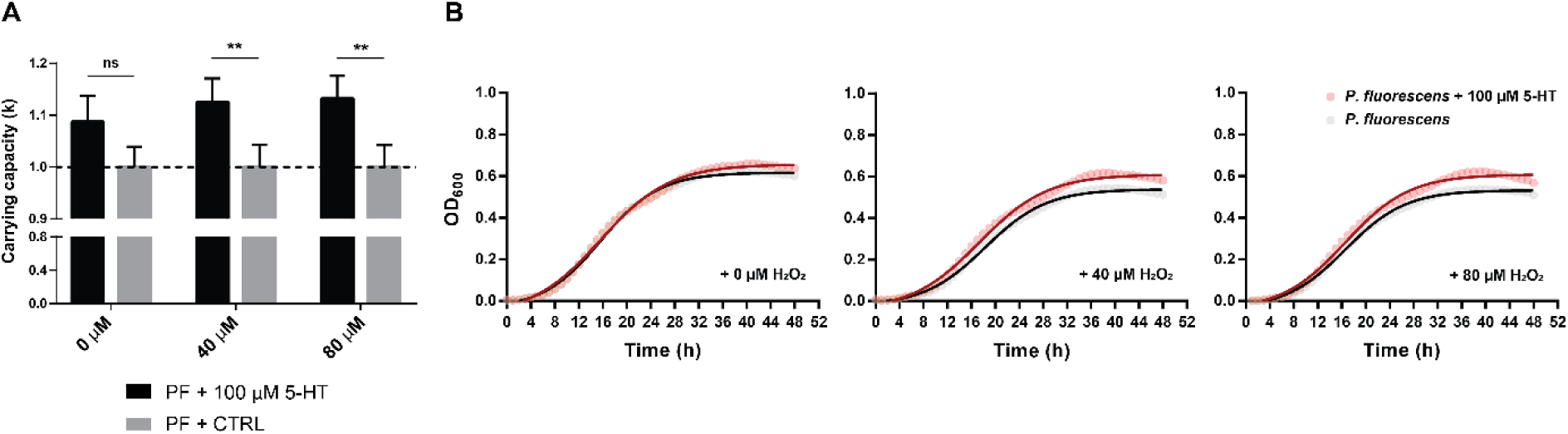
Serotonin metabolism protects *P. fluorescens* from oxidative stress. (A) Quantification of carrying capacity k (a measure of maximum OD600) after P. fluorescens stimulation with or without 100 µM serotonin and in the presence (40 and 80 µM H2O2) or absence (no addition of H2O2) of oxidative stress. Data represent average ± SEM. ns = not significant. Statistical significance was calculated using two-tailed unpaired t-tests. (B) Representative growth curves of P. fluorescens stimulated with or without 100 µM serotonin and in the presence (40 and 80 µM H2O2) or absence (no addition of H2O2) of oxidative stress.

## Discussion

In this study, we demonstrated the ability of *P. fluorescens* to metabolize serotonin into several metabolic products, including 5-hydroxytryptophol, 5-HIAA, and N-acetylserotonin, with 5-HIAA being the most abundant product (**Figure 1**). Our RNA-seq and proteomics analyses revealed that serotonin exposure significantly upregulates several functional and metabolic pathways, particularly those involved in oxidative stress defense, such as amino acid catabolism (phenylalanine, tyrosine, and arginine degradation pathways) (**Figure 2B,D**). Furthermore, serotonin was shown to protect *P. fluorescens* from oxidative stress induced by H_2_O_2_ exposure (**Figure 3**), supporting its role as an antioxidant. These findings suggest that serotonin not only serves as a metabolic substrate for *P. fluorescens*, but also plays a crucial role in protecting the bacterium from environmental stressors, including the gut.

Recent studies have shown that serotonin can enhance gut bacterial fitness by upregulating genes related to growth and survival (Fung et al. 2019). Although our study, along with others (Biaggini et al. 2015), did not observe any direct impact of serotonin on *P. fluorescens* growth (**Figure 1G**), our RNA-seq analysis revealed significant upregulation of several functional pathways upon serotonin exposure. These include CoA-transferase activity, polysaccharide transport, the beta-ketoadipate pathway, and transcription regulation (**Figure 2B**). The beta-ketoadipate pathway, which is commonly found in aerobic bacteria like *Pseudomonas*, plays a key role in the degradation of aromatic compounds such as benzoate and catechol (D. Li et al. 2010). These compounds serve as vital carbon and energy sources for many microorganisms, feeding into the tricarboxylic acid (TCA) cycle (Suvorova and Gelfand 2019). Furthermore, we observed significant upregulation of transcriptional regulators, including *gntR* and *cra*, both of which play roles in carbon metabolism *in P. aeruginosa* and may suggest an adaptive response in *P. fluorescens.* GntR has been implicated in glucose metabolism regulation in *P. aeruginosa* (Daddaoua et al. 2017), while *cra* encodes the catabolite repressor/activator Cra, which modulates carbohydrate uptake via the phosphoenolpyruvate: phosphotransferase system (PTS) in *P. putida* (Chavarría et al. 2013). Our proteomics data further supported the RNA-seq findings, showing upregulation of pathways related to energy production and conversion, carbohydrate transport and metabolism, and amino acid transport and metabolism.

In disease states such as celiac disease, bacteria are continuously exposed to oxidative stress, whether endogenously produced via respiration or exogenously induced by the host immune system (da Cruz Nizer et al. 2021). To survive these conditions, bacteria have evolved adaptive mechanisms to mitigate oxidative damage (da Cruz Nizer et al. 2021). Among these, the degradation pathways of phenylalanine, tyrosine, and arginine have been identified as crucial for cellular protection against oxidative stress (Id et al. 2019; Ipson and Fisher 2016; Tiwari et al. 2018). Under oxidative stress, hydroxyl radicals can oxidize the benzyl ring of phenylalanine, leading to the formation of abnormal tyrosine isomers, meta-tyrosine and ortho-tyrosine, which can be toxic to cells (Ipson and Fisher 2016). One key defense strategy involves their elimination through metabolic degradation pathways, including the phenylalanine and tyrosine catabolic pathway, which is linked to the production of pyomelanin — a secondary metabolite known for its protective role against oxidative stress in *Pseudomonas* (Turick et al. 2010). Similarly, arginine associated pathway was shown to be one of the early adaptive responses to oxidative stress in *Mycobacterium tuberculosis* (Tiwari et al. 2018). Similarly, the arginine-associated pathway plays an early adaptive role in oxidative stress responses, as demonstrated in *Mycobacterium tuberculosis*.

Serotonin itself, along with its degradation product N-acetylserotonin, has been reported as an antioxidant across different biological systems (Wan et al. 2018; Azouzi et al. 2017; Gonçalves et al. 2022; Liang et al. 2020; Kang et al. 2009). In plants, serotonin accumulation under stress conditions is linked to reactive oxygen species (ROS) scavenging and modulation of stress responses (Kang et al. 2009). *In vitro*, serotonin has been shown to protect lipid membranes from oxidation (Azouzi et al. 2017). Together, these findings support our conclusion that serotonin catabolism in *P. fluorescens* triggers metabolic responses, including the phenylalanine, tyrosine, and arginine degradation pathways (**Figure 2**), to combat oxidative stress. This protective effect may enhance bacterial survival against host oxidative immune responses, particularly in disease contexts such as celiac disease.

## Conclusion

This study demonstrates that *P. fluorescens* MFY63 efficiently metabolizes serotonin, primarily yielding 5-HIAA. Transcriptomic and proteomic analyses revealed that serotonin exposure significantly upregulates amino acid catabolic pathways, including phenylalanine, tyrosine, and arginine degradation—pathways known to mitigate oxidative stress. Additionally, serotonin metabolism enhanced bacterial resistance to oxidative stress, suggesting a protective role. These findings highlight a potential mechanism by which *P. fluorescens* adapts to oxidative environments, potentially promoting its persistence in the gastrointestinal tract. While the direct impact on celiac disease pathophysiology remains to be determined, our data indicate that serotonin metabolism may provide *P. fluorescens* with a survival advantage in the intestinal environment. Future studies incorporating host-microbe interactions and patient-derived samples will be essential to clarify the clinical relevance of this interaction.

## Materials and Methods

### Bacterial strains and growth conditions

To screen for the conversion capacity of gut bacteria species, 14 aerobic and 18 anaerobic strains were tested in **Supplementary Table 1.** Strains were grown either aerobically at 37 °C with continuous agitation or anaerobically (10% H_2_, 10% CO_2_, 80% N_2_) in a Don Whitley Scientific DG250 Workstation (LA Biosystems, Waalwijk, the Netherlands) at 37 °C in a fresh Enriched Beef Broth (EBB) (**Supplementary Table 11**). Anaerobic bacteria were inoculated from −80 °C stocks and grown overnight. Before the experiment, cultures were diluted 1:100 in fresh medium from overnight liquid cultures. Serotonin hydrochloride (5-HT; H9523; Sigma) was supplemented during the lag phase of a bacterial growth. The bacterial cultures were incubated at 37 °C with continuous agitation for 24 h in Erlenmeyer flasks. Samples (2mL) were collected after 24 h of incubation and supernatant was taken after centrifugation at 20,000 × g for 20 min at 4 °C.

*P. fluorescens* MFY63 was first restreaked from −80 °C stocks onto LB (Luria broth) plate and incubated at 37 °C for at least 24 h. Single colony was then inoculated into an EBB and grown overnight at 37 °C with continuous agitation. To decipher the serotonin catabolism and further molecular analysis of *P. fluorescens* MFY63, the grown overnight culture of *P. fluorescens* MFY63 was diluted 1:100 to a fresh EBB medium. To each of the suspensions, 100 µM 5-HT or sterile MilliQ-filtered water was supplemented. Growth was followed by measuring the optical density (OD) at 600 nm in a spectrophotometer (UV1600PC, VWR International, Leuven, Belgium). The bacterial cultures were incubated at 37 °C with continuous agitation for 24 h in Erlenmeyer flasks. Samples (2mL) were collected after 6, 8, 10, 12 and 24 h of incubation. The supernatant and bacterial pellets were obtained after centrifugation at 20,000 × g for 20 min at 4 °C and stored in −80 °C until further analysis. The samples were used further for LC-MS/MS and RNA-seq analysis.

### Extraction of metabolites and LC-MS/MS

A total of 1 mL of collected culture supernatant was mixed with 2 mL ethylacetate (EtOAc), and incubated overnight at 4 °C. Then, 1 mL of the upper organic layer was collected, and another 1 mL fresh EtOAc was added to the supernatant/EtOAc mixture and mixed. After a few hours of incubation, again 1 mL of the upper layer was collected and added to the first collection fraction. The solvent was evaporated from the collected extract under nitrogen flow. For LC-MS/MS analyses, the extracts were redissolved in 1 mL methanol. The *N*-acetylserotonin standard purchased from Sanbio B.V. was added in methanol to a final concentration of 10 µg/mL.

LC-MS/MS acquisition was performed using Shimadzu Nexera X2 UHPLC system, with attached PDA, coupled to Shimadzu 9030 QTOF mass spectrometer, equipped with a standard ESI source unit, in which a calibrant delivery system (CDS) is installed. 2 µL of the extracts were injected into a Waters Acquity HSS C18 column (1.8 μm, 100 Å, 2.1 × 100 mm). The column was maintained at 30 °C, and run at a flow rate of 0.5 mL/min, using 0.1% formic acid in H_2_O as solvent A, and 0.1% formic acid in acetonitrile as solvent B. A gradient was employed for chromatographic separation starting at 5% B for 1 min, then 5–85% B for 9 min, 85–100% B for 1 min, and finally held at 100% B for 3 min. The column was re-equilibrated to 5% B for 3 min before the next run was started. The LC flow was switched to the waste the first 0.5 min, then to the MS for 13.5 min, then back to the waste to the end of the run. The PDA acquisition was performed in the range 200–600 nm, at 4.2 Hz, with 1.2 nm slit width. The flow cell was maintained at 40 °C.

All the samples were analyzed in positive polarity, using data dependent acquisition mode. In this regard, full scan MS spectra (m/z 100–1700, scan rate 10 Hz, ID enabled) were followed by two data dependent MS/MS spectra (m/z 100–1700, scan rate 10 Hz, ID disabled) for the two most intense ions per scan. The ions were selected when they reach an intensity threshold of 1500, isolated at the tuning file Q1 resolution, fragmented using collision induced dissociation (CID) with fixed collision energy (CE 20 eV), and excluded for 1 s before being re-selected for fragmentation. The parameters used for the ESI source were interface voltage 4 kV, interface temperature 300 °C, nebulizing gas flow 3 L/min, and drying gas flow 10 L/min. The parameters used for the CDS probe were interface voltage 4.5 kV, and nebulizing gas flow 1 L/min.

All raw data is available at MassIVE under the following dataset identifier: MSV000097220

### Metabolomics data analysis

Raw data obtained from LC-MS analysis were converted to mzXML centroid files using Shimadzu LabSolutions Postrun Analysis. The files were imported into Mzmine 2.53 for data processing (Pluskal et al. 2010). Unless stated otherwise, m/z tolerance was set to 0.002 m/z or 15.0 ppm, RT tolerance was set to 0.05 min, noise level was set to 2.0E2 and the minimum absolute intensity was set to 5.0E2. Mass ion peaks were detected (positive polarity, mass detector: centroid) and their chromatograms were built using ADAP chromatogram builder (minimum group size in number of scans: 5; group intensity threshold: 2.0E2). The detected peaks were smoothed (filter width: 9), and the chromatograms were deconvoluted (algorithm: local minimum search; chromatographic threshold: 90 %; search minimum in RT range: 0.05; minimum relative height: 1 %; minimum ratio of peak top/edge: 2; peak duration: 0.03–3.00 min). The detected peaks were deisotoped (monotonic shape; maximum charge: 2; representative isotope: most intense). Peak lists from different extracts were aligned (weight for RT = weight for m/z = 20; compare isotopic pattern with a minimum score of 50%). Missing peaks detected in at least one of the samples were filled with the gap filling algorithm (RT tolerance: 0.1 min). Duplicate peaks were filtered. Artifacts caused by detector ringing were removed (m/z tolerance: 1.0 m/z or 1000.0 ppm). Only features with RT 0.5–10 min were kept. The aligned peaks were exported to a MetaboAnalyst file.

In Excel, features that were not present with an intensity higher than 3000 in at least two samples out of three from the same condition (each condition was measured in triplicates) were removed from the file. Additionally, all features that originate from the culture medium were removed by retaining only features with an average peak intensity of at least 20 times greater in the bacterial extracts than in the culture medium extracts. The resulting peak list was uploaded to MetaboAnalyst (Xia et al. 2009) for statistical analysis. Log transformation with pareto scaling was applied to the data. Differences with a two-fold change and an FDR-adjusted *p*-value < 0.1 were considered statistically significant (unless otherwise stated).

### Collection of the small intestinal content and incubation experiments

Small intestinal content (SIC) was collected from five male WTG rats (retired breeders) that had been used in a separate study (Waclawiková et al., 2021). Luminal contents were harvested from the entire jejunum by gentle pressing, snap-frozen in liquid nitrogen, and stored at −80°C until use in incubation experiments. For an incubation experiments, luminal contents from the jejunum of wild-type Groningen rats (n = 5) were suspended in EBB (5% w/v) containing 100 µM 5-HT and were spiked with either 0%, 1% or 10% of grown bacterial culture of *P. fluorescens* and incubated aerobically with continuous agitation for 24 h at 37 °C prior to the HPLC-ED analysis.

### HPLC-ED analysis and sample preparation

A volume of 1 mL of ice-cold methanol was added to 0.25 mL bacterial cell suspensions from 5-HT incubation experiments with SIC samples and with *P. fluorescens*, and stored at −20 °C until further use. Cell and protein precipitates were removed by centrifugation at 20,000 × g for 10 min at 4 °C. Supernatant was transferred to a new tube, and the methanol fraction was evaporated in a Savant speed-vacuum dryer (SPD131, Fisher Scientific, Landsmeer, the Netherlands) at 60 °C for 1 h 15 min. The aqueous fraction was reconstituted to 1 mL with 0.7% HClO_4_. Samples were filtered and injected into the HPLC system (Waters 2695 Alliance Separations module, Milford, Massachusetts, USA; Dionex ED40 electrochemical detector, Dionex, Sunnyvale, USA, with a glassy carbon working electrode (DC amperometry at 1.0 V, with Ag/AgCl as reference electrode). Samples were analyzed on a C18 column (Kinetex 5 μm, C18 100 Å, 250 × 4.6 mm, Phenomenex, Utrecht, the Netherlands) using a gradient of water/methanol with 0.1% formic acid (0 to 10 min, 95% to 80% H_2_O; 10 to 20 min, 80% to 5% H_2_O; 20 to 23 min, 5% H_2_O; 23 to 31 min, 95% H_2_O). Data recording and analysis were performed using Chromeleon software (version 6.8 SR13) and GraphPad Prism 7.

### Genomic DNA extraction

Prior to genomic DNA (gDNA) isolation, *P. fluorescens* MFY63 was cultured in EBB overnight at 37 °C and cells were harvested in the stationary growth. gDNA was extracted from grown *P. fluorescens* MFY63 culture using GenElute^TM^ Bacterial Genomic DNA Kit (NA2110; Sigma Aldrich) and manufacturer’s protocol for gram-negative bacteria was followed.

### Genome sequencing

Whole genome sequencing was performed on a MGISEQ-2000 system at BGI Tech Solutions (Hong Kong) and 150 bp reads were generated with a library insert size of 350 bp. Briefly, 1μg genomic DNA was randomly fragmented using a g-TUBE device (Covaris, Inc) according to the manufactures instructions. The fragmented DNA was size selected using magnetic beads to an average of 200-400bp and selected fragments were end-repaired, 3’ adenylated, adapter-ligated and PCR amplified. The double stranded PCR product was heat denatured and circularized.

After sequencing reads were quality controlled using FastQC v0.11.9 (Andrews 2010) was and low-quality reads were removed with Trimmomatic v0.38 (Bolger, Lohse, and Usadel 2014). The reads were then *de novo* assembled using SPAdes v3.11.1 with default parameters (Bankevich et al. 2012). Finally the genome sequence was annotated using Prokka under default settings (Seemann 2014). The genome is available in GenBank under JAGKIG000000000.

### RNA extraction

Bacterial pellets were resuspended (by vortexing) in 100 µL of Buffer A [10 mM Tris pH 8.0; 100 mM NaCl], supplemented with RNase inhibitor (EO1861F, Applied Biosystems, Thermo Fisher Scientific). After resuspension, 25 - 100 µL of Buffer B [50 mM EDTA; 120 mM Tris pH 8.0], supplemented with lysozyme (100 ug/mL final), was added to each suspension. Suspensions were tilted 5-10 times to ensure homogenous mixing and then incubated 1 min at room temperature. Subsequently, 125 −500 µL of Buffer C [0.5% Tween-20; 0.4% sodium deoxycholate; 2 M NaCl; 10 mM EDTA] was added, suspensions were mixed 5-10 times by tilting and incubated for 5 min at room temperature. Next, 100 µL of each suspensions was taken and transferred to a new tube containing 1 mL ice-cold TRIzol (15596018, Ambion, Thermo Fisher Scientific) and samples were vortexed thoroughly. If the nucleoid was visible (or the mixture appeared viscous), sample was incubated at 56 °C for 5 min. TRIzol RNA extraction protocol was followed immediately afterwards. Samples were allowed to stand for 5 min at room temperature and then 200 µL chloroform (without additives) was added. Samples were shaken for 15 s and then incubated for 5-10 min at room temperature. After incubation, samples were centrifuged for 15 min at 12,000 x g at 4 °C. Colorless upper aqueous phase (containing RNA) was transferred to a new tube containing 500 µL isopropanol and suspensions were mixed by pipetting and incubated for 5-10 min at room temperature. Next, samples were centrifuged for 10 min at 12,000 x g at 4 °C. The RNA precipitate formed a visible pellet on the side and the bottom of the tube. The supernatants were removed and 1 mL of 75% absolute ethanol was added. Samples were vortexed and centrifuged for 5 min at 12,000 x g at 4 °C. After centrifugation, the supernatants were carefully removed and RNA pellets were dried for 5 min by air drying. Extensive drying of the RNA samples was avoided to ensure sufficient solubility of the samples. The RNA pellets were resuspended in 20 µL of DEPC-treated water heated at 60 °C prior dissolution. RNA concentration and quality was checked on the Nanodrop spectrophotometer (Nanodrop 2000, Isogen, De Meern, The Netherlands) and total RNA was loaded on 1% bleach gel (500 µL of 6% sodium hypochlorite (bleach) was added to 50 mL 1% agarose gel; 1 µL of total RNA was loaded on the gel and the electrophoresis was run for 30 min at 100 volt). The rest of the RNA samples was stored at −80 °C until further procedure.

### RNA library preparation, RNA sequencing and RNA-seq analysis

The Zymo-Seq RiboFree™ Total RNA Library Kit (R3000, Zymo Research, USA) was used to prepare the library preps for Illumina sequencing using 3 µg of the extracted total RNA. Samples were sequenced on the Illumina NexSeq 500 to generate 75 bases single end reads (75SE) with an average read-depth of 12M reads per sample. All raw reads are available at SRA under PRJNA930327. The quality of the resulting fastq reads we checked using FastQC v0.11.9 (Andrews 2010) and mapped on the reference genome of *P. fluorescens* MFY63 (GenBank: JAGKIG000000000) using Bowtie2 v2.4.2 using default settings (Langmead and Salzberg 2012). Resulting SAM files were converted to BAM using SAMtools 1.11 and featuresCounts 2.0.1 was used to get the gene counts (H. Li et al. 2009; Liao, Smyth, and Shi 2014). The T-REx webserver was used to perform statistical analysis and determine differential gene expressions (DGE) (de Jong et al. 2015) and subsequently, gene set enrichment analysis was done for functional analysis using the GSEA-Pro web server (http://gseapro.molgenrug.nl) or KEGG database (Kanehisa et al. 2017). Heatmaps containing differentially expressed genes with FDR < 0.05 were generated using the ComplexHeatmap package for R (Gu, Eils, and Schlesner 2016).

### Proteomics Sample preparation

Samples were prepared essentially as described (Gubbens et al. 2012). Briefly, after incubation with 100 µM 5-HT, bacterial culture was pelleted and lysed in 100 µL lysis buffer (4% SDS, 100 mM Tris-HCl pH 7.6, 50 mM EDTA) and disrupted by sonication. Total protein was precipitated using the chloroform-methanol method (Wessel and Flügge 1984), and the proteins dissolved in 0.1% RapiGest SF surfactant (Waters) at 95 °C. The protein concentration was measured at this step using BCA method. Protein samples were then reduced by adding 5 mM DTT and incubate at 60 °C for 30 min, followed by thiol group protection with 21.6 mM iodoacetamide incubation at room temperature in dark for 30 min. Then 0.1 μg trypsin (recombinant, proteomics grade, Roche) per 10 μg protein was added, and samples were digested at 37 °C overnight. After digestion, trifluoroacetic acid was added to 0.5% and samples were incubated at 37 °C for 30 min followed by centrifugation to degrade and remove RapiGest SF. Peptide solution containing 6 μg peptide was then cleaned and desalted using STAGE-Tips (Rappsilber, Mann, and Ishihama 2007). Briefly, 6 μg of peptide was loaded on a conditioned StageTip with 2 pieces of 1 mm diameter C18 disk (Empore, product number 2215), washed twice with 0.5 % formic acid solution, and eluted with elution solution (80% acetonitrile, 0.5% formic acid). The acetonitrile was then evaporated in a SpeedVac. Final peptide concentration was adjusted to 40 ng·μL^-1^ using sample solution (3% acetonitrile, 0.5% formic acid) for analysis.

### Proteomics LC-MS/MS Measurement

The desalted peptides solution was separated on an UltiMate 3000 RSLCnano system set in a trap-elute configuration with a nanoEase M/Z Symmetry C18 100 Å, 5 µm, 180 µm × 20 mm (Waters) trap column for peptide loading/retention and nanoEase M/Z HSS C18 T3 100 Å, 1.8 µm, 75 µm × 250 mm (Waters) analytical column for peptide separation. Mobile phase A composed of 0.1% formic acid (FA) in ULC-MS grade H2O (Biosolve), while mobile phase B composed of 0.1% FA, 10% H2O in ULC-MS grade acetonitrile (ACN, Biosolve). The flow gradients used for analysis was a shallow 113 min gradient of mobile phase A and B controlled by a flow sensor at 0.3 µL·min^-1^. The gradient was programmed with linear increment from 1% to 5% B from 0 to 2 min, 5% to 13% B from 2 to 63 min, 13% to 22% B from 63 to 85 min, 22% to 40% B from 85 to 104 min, 90% at 105 min and kept at 90% to 113 min.

The eluent was introduced by electro-spray ionisation (ESI) via the nanoESI source (Thermo) to QExactive HF (Thermo Scientific). The QExactive HF was operated in positive mode with data dependent acquisition. The MS survey scan was set with mass range m/z 350-1400 at 120,000 resolution. For individual peaks, the data dependent intensity threshold of 2.0 × 104 was applied for triggering an MS/MS event, isotope exclusion on and dynamic exclusion was 20 s. Unassigned, +1 and charges >+8 were excluded with peptide match mode preferred. For MS/MS events, the loop count was set to 10, isolation window at m/z 1.6, resolution at 15,000, fixed first mass of m/z 120, and normalised collision energy (NCE) at 28 eV.

All raw data is available at MassIVE under the following dataset identifier:

### Proteomics data analysis

Raw proteomics data was processed using MaxQuant (version 2.1.0.0) (Cox and Mann 2008; Tyanova, Temu, and Cox 2016) with label free quantification (LFQ) enabled. The result was further processed using MSstats (version 4.2.0) (Choi et al. 2014) to calculate the differentially presented proteins. To assess the differentially abundant proteins of *P. fluorescens* grown in presence of 100 µM 5-HT compared to the control group, the volcano plots were build using R, where significant features were filtered by an FDR-adjusted p-value < 0.05, and two-fold change.

### Oxidative stress assay

P. fluorescens MFY63 was first restreaked from −80 °C stocks onto LB (Luria broth) plate and incubated at 37 °C for at least 24 h. Single colony was then inoculated into a fresh EBB medium and grown overnight at 37 °C with continuous agitation. Before the experiment, cultures were diluted 1:100 in fresh EBB medium from overnight culture. To each of the suspensions, 100 µM 5-HT or sterile MilliQ-filtered water together with or without 40 µM or 80 µM H_2_O_2_ was supplemented. Bacterial growth was monitored for 48 h at 600 nm (OD600) with shaking for 5s every 10 min using a BioTek ELx808 microplate reader (BioTek, Winooski, USA). Growth curves were fitted using the growthcurver R package (v0.3.1) (Sprouffske and Wagner 2016).

### Statistics

Results are expressed as means from two to five biological replicates with the corresponding standard deviations. The growth curves were fitted using the growthcurver R package (v0.3.1) (Sprouffske and Wagner 2016).

## Acknowledgments

We thank Prof. Dr. Marc G.J. Feuilloley, Department of the Laboratory of Microbiology Signals and Microenvironment, University of Rouen, France, for providing us with *Pseudomonas* strains (Chapalain et al., 2008); Dr. Hermie J. M. Harmsen, Department of Medical Microbiology, University Medical Centre, Groningen, The Netherlands, for providing us with gut bacterial strains used in the screening study; Dr. Elisa Brambilla, Max Planck Institute for Evolutionary Biology, Ploen, Germany, for providing us with *P. fluorescens* SBW25 strain used in the screening study; Dr. Emma Allen-Vercoe, University of Guelph, Ontario, Canada, for providing us with *F. nucleatum* subsp. *animalis* EAVG_002 used in the screening study; Dr. Marten Exterkate for assistance with running and analyzing the samples on the LC-MS; Warner Hoornenborg, Department of Molecular Neurobiology, University of Groningen, the Netherlands, for assistance with the collection of rat small intestinal content; Dr. Danny Incarnato, Department of Molecular Genetics, University of Groningen, the Netherlands, for help with setting up the RNA-seq study.

## Financial Disclosure

S.E.A. is supported by Rosalind Franklin Fellowships, co-funded by the European Union and University of Groningen. The funders had no role in study design, data collection and analysis, decision to publish, or preparation of the manuscript.

## Author Contributions

B.W. and S.E.A. conceptualized and designed the study. B.W. performed the experiments and analysis of the data. M.S. assisted with bioinformatics and analysis of the data. S.T. assisted with 5-HT degradation overtime experiment. A.d.J. performed genome sequencing analysis and assisted with analysis of RNA-seq data. S.E.A. assisted with analysis of the data. B.W. wrote the original manuscript D.I. contributed to restructuring and editing the manuscript that was reviewed by M.S., S.T., A.d.J., and S.E.A. Funding was acquired by S.E.A.

## Competing Interests Statement

The authors have declared that no competing interests exist.

## Notes

### Competing Interest Statement

The authors have declared no competing interest.

